# Highly pathogenic avian influenza causes mass mortality in Sandwich tern (*Thalasseus sandvicensis*) breeding colonies across northwestern Europe

**DOI:** 10.1101/2023.05.12.540367

**Authors:** Ulrich Knief, Thomas Bregnballe, Ibrahim Alfarwi, Mónika Ballmann, Allix Brenninkmeijer, Szymon Bzoma, Antoine Chabrolle, Jannis Dimmlich, Elias Engel, Ruben Fijn, Kim Fischer, Bernd Hälterlein, Matthias Haupt, Veit Hennig, Christof Herrmann, Ronald in ‘t Veld, Elisabeth Kirchhoff, Mikael Kristersson, Susanne Kühn, Kjell Larsson, Rolf Larsson, Neil Lawton, Mardik Leopold, Sander Lilipaly, Leigh Lock, Régis Marty, Hans Matheve, Włodzimierz Meissner, Paul Morisson, Stephen Newton, Patrik Olofsson, Florian Packmor, Kjeld T. Pedersen, Chris Redfern, Francesco Scarton, Fred Schenk, Olivier Scher, Lorenzo Serra, Julian Smith, Wez Smith, Jacob Sterup, Eric Stienen, Viola Strassner, Roberto G. Valle, Rob S. A. van Bemmelen, Jan Veen, Muriel Vervaeke, Ewan Weston, Monika Wojcieszek, Wouter Courtens

## Abstract

In 2022, highly pathogenic avian influenza (HPAI) A(H5N1) virus clade 2.3.4.4b became enzootic and caused mass mortality in Sandwich terns and other seabird species across northwestern Europe. We present data on characteristics of the spread of the virus between breeding colonies and the number of dead adult Sandwich terns recorded at breeding sites throughout northwestern Europe. Within two months after the first mortalities were reported, in total 20,531 adult Sandwich terns were found dead, which is >17% of the total northwestern European breeding population. Losses are likely higher, as we expect that many victims were not found (mortality rate might be up to 74% of the breeding population). Inside the colonies almost all chicks died. After the peak of the outbreak, in a colony established by late breeders, 25.7% of adults showed immunity against HPAI subtype H5. Removal of carcasses helped in reducing the spread of the disease and consequently total mortality. More research on the sources and modes of transmission, incubation times, effective containment and immunity is urgently needed to combat this major threat for colonial seabirds.

## Introduction

Highly pathogenic avian influenza (HPAI) A(H5N1) clade 2.3.4.4b caused the largest and most severe global epizootic event in poultry, captive and wild birds witnessed to date (*1-4*). Until 2021, avian influenza viruses in Europe occurred seasonally, with the majority of infections being reported between autumn and spring and no recognized infections over the summer months (*1, 5*). While in winter 2020/21 subtype HPAI H5N8 dominated the epizootic and led to severe mortality among barnacle (*Branta leucopsis*) and greylag geese (*Anser anser*; *6-8*), the subtype HPAI H5N1 clade 2.3.4.4b dominated in wild birds across Europe from late spring 2021 onward. Infections were detected sporadically throughout the summer and early autumn of 2021, causing mass mortality in great cormorants (*Phalacrocorax carbo*) in Estonia in May 2021 (*9*) and in great skuas (*Stercorarius skua*) in the United Kingdom in July 2021 (*10*). However, in autumn and winter 2021/22, Europe witnessed an unprecedented outbreak of HPAI H5N1 in both wild birds and poultry (*1*). The virus was detected in at least 62 wild bird species of which 17 were waterfowl and 12 were raptor species (*5*). The most severe cases were the losses of more than 4,500 barnacle geese of the Svalbard breeding population that died on the Solway Firth (UK) and the associated decline of the wintering population by 38% (*11*), the death of 5,000 common cranes (*Grus grus*) in northern Israel and of 2,000 Dalmatian pelicans (*Pelecanus crispus*) in Greece (*2, 12, 13*).

By spring 2022, HPAI H5N1 clade 2.3.4.4b had become enzootic, meaning that it did not disappear during the summer months, and caused the largest ever recorded outbreak in wild birds and poultry across Europe (*14, 15*), and a subsequent spread to the Americas (including previously spared South America; *2, 4, 16-18*). In Europe, almost 2,500 outbreaks among poultry with 47.7 million birds culled between October 2021 and September 2022 signified an enormous economic loss and pressure on the food chain (*19*). Between March and September 2022, the virus was detected in 80 wild avian species (*15, 19*), with some colonially nesting seabirds being affected the most. Great skuas, northern gannets (*Morus bassanus*), great cormorants, common terns (*Sterna hirundo*) and Sandwich terns (*Thalasseus sandvicensis*) all experienced major die-offs (*19-23*).

Sandwich terns are highly gregarious seabirds, with the majority of the European population breeding in a small number of large coastal colonies (*24*), that are annually and often comprehensively monitored (cf. Annex I of the EC Birds Directive and thus a conservation priority). This allowed following the spread and the impact of HPAI in this wild avian species closely. The European Sandwich tern breeding population consists of three flyway populations, of which the northwestern European population was worst affected by HPAI in 2022. Its size is estimated at 45,600–55,300 breeding pairs (BP) or 170,000 individuals (*25*).

Sandwich terns nest at densities of up to 7 BP/m^2^ with an average distance between nests of 30–35 cm (*26; own observations*). Contrary to other European tern species, adults defecate around the nest while breeding, to an extent that vegetation growth is hampered by toxic amounts of nutrients (*26-29*). Adults are highly mobile, moving within and between colonies, both within and across breeding seasons (*30-33*). Viral transmission via the fecal-oral route probably predominates among wild birds (*34*). The Sandwich tern’s behavior promotes airborne and direct-contact transmission routes and may put the species at high risk for a HPAI epizootic once the virus has entered the population (*35*).

Sandwich terns are long-lived, with an adult annual survival rate of 0.92 (*36*) and a recorded maximum age of 30 years (*37*). Their reproductive output is usually low (<1 fledgling per BP and year; *38*), which makes adult survival of paramount importance in population dynamics (*35*). Thus, we here focus mainly on the impact of HPAI on adult mortality, describe the spatiotemporal patterns of infection across all Sandwich tern colonies in northwestern Europe and evaluate containment, preventive and mitigating measures that were pursued during the breeding season in 2022.

## Results and Discussion

We collected data from 67 colonies that in total harbored 63,116 BP, which agrees with the estimated entire known northwestern European breeding population (*25, 39*). HPAI was confirmed in 39 out of 65 colonies (60%; **Fig. 1**), comprising 74% of the breeding pairs (*N* = 46,701 BP). All confirmed infections were caused by H5N1. In total, 16,873 adult Sandwich terns were found dead inside or near the affected colonies (18.1% and 13.4% of the breeding birds in affected and all colonies, respectively). For a subset of regions, dead adults were also counted along the shorelines (BE: 59, NL: 1,600, DE NI: 598, DE SH: 196 dead birds) that likely died there or at sea while foraging. Using these data, we estimated that an additional 3,658 adult dead Sandwich terns stranded across northwestern Europe outside the colonies (see Methods). Thus, at least 16.3% (= [16,873 + 3,658] / [2 × 63,116]) of adult Sandwich terns of the entire northwestern European breeding population died during the breeding season of 2022. This figure is an underestimate of the actual rate (see below), but nevertheless an order of magnitude higher than the expected “natural” (that is pre-HPAI) rate of ∼1.3% (= 8% yearly mortality / 12 months × 2 months of breeding season; cf. *36*). Some birds certainly rebred after losing their first clutch (*40*; *N* = 3,484 BP in colonies formed late in the season), leading to a mortality rate >17%. As many birds undoubtedly were lost at sea, beached without being reported, or died on land away from colonies (*41*), the actual mortality rate must have been substantially higher (in the range of 17–74%, if all birds in affected colonies died). Supporting evidence comes from standardized seawatching observations conducted in the Netherlands during autumn migration, where passing rates of Sandwich terns were 70–80% lower compared to previous years (*21*), including a much reduced percentage of first-year birds at a major northwestern European roost in autumn 2022 (percentages in 2020/21 = 37.2% vs 6.9% in 2022; estimated contrast = 31.0% [95% confidence interval CI = 25.2%–36.8%], *P* < 0.001; **Fig. S1**), because most of the chicks died in affected colonies, either because of the HPAI infection itself or because at least one of their parents died in the outbreak.

**Fig. 1.**
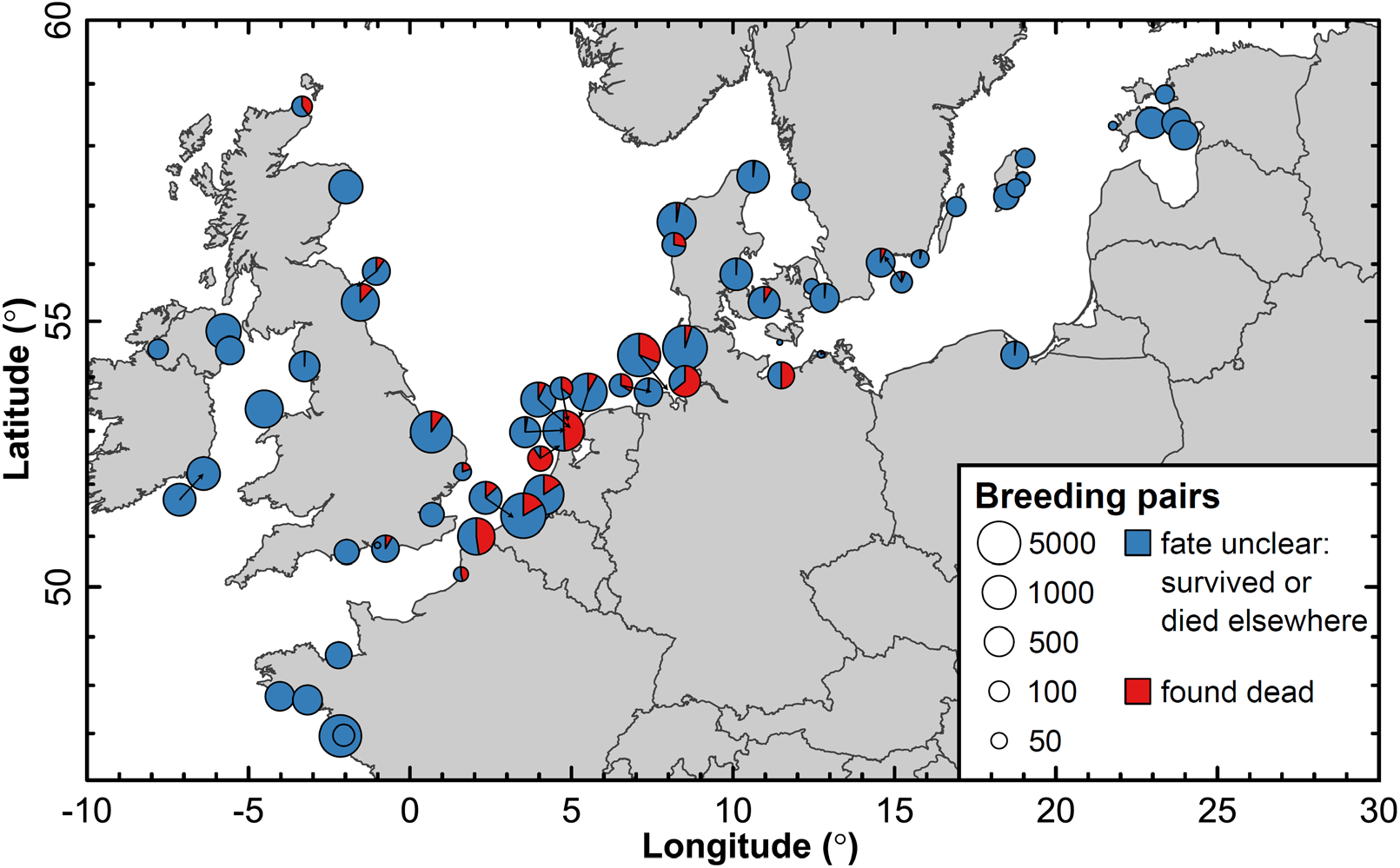
Distribution of Sandwich tern colonies and death rates of adult breeding birds due to HPAI in northwestern Europe. Dot size reflects colony size and pie charts represent the percentages of adults with an unknown survival fate (that is, either survived or died elsewhere; blue) and found dead (red) in a colony.

The HPAI outbreak did not reach all Sandwich tern colonies in northwestern Europe. The risk of a colony being infected increased with its size (*β* = 0.74 [95% CI = 0.019–1.45], *P* = 0.044) and with its position relative to the center of the species’ breeding range in northwestern Europe (*β* = 4.86 [95% CI = 1.33–8.38], *P* = 0.007; **Fig. 1**). Specifically, colonies towards the periphery in the Baltic Sea (eastern SE, EE), in the Irish Sea (western UK, IE) and the Atlantic (southwestern FR) were not affected, whereas the large colonies along the shores of the southern North Sea (northern FR, BE, NL, western DE, southeastern UK) suffered major losses. Overall, this suggests that adult Sandwich terns transmitted the virus between colonies, as larger and more centrally located colonies are expected to attract more visitors than smaller, more peripheral ones. Indeed, ring recovery data suggests a high visiting rate of birds from other colonies in a large, central colony (*30*), and that British and Irish colonies are partially isolated from the continental population (*42, 43*).

HPAI was likely introduced into the Sandwich tern breeding population in 2022 at least three times independently, as there were two simultaneous outbreaks occurring end of April/beginning of May in eastern Germany and in northern Scotland and a third outbreak middle of May in northern France (**Fig. 2**). This was supported by genetic analyses of whole viral genomes (*21, 22*): the eastern German outbreak seemed to be caused by a single virus variant (cluster 2 in *22*), whereas the French and Dutch outbreaks were caused by two variants of HPAI H5N1 clade 2.3.3.4.b (clusters 2 and 3 in *21, 22*). The variants causing the outbreak in northern Scotland were not determined.

**Fig. 2.**
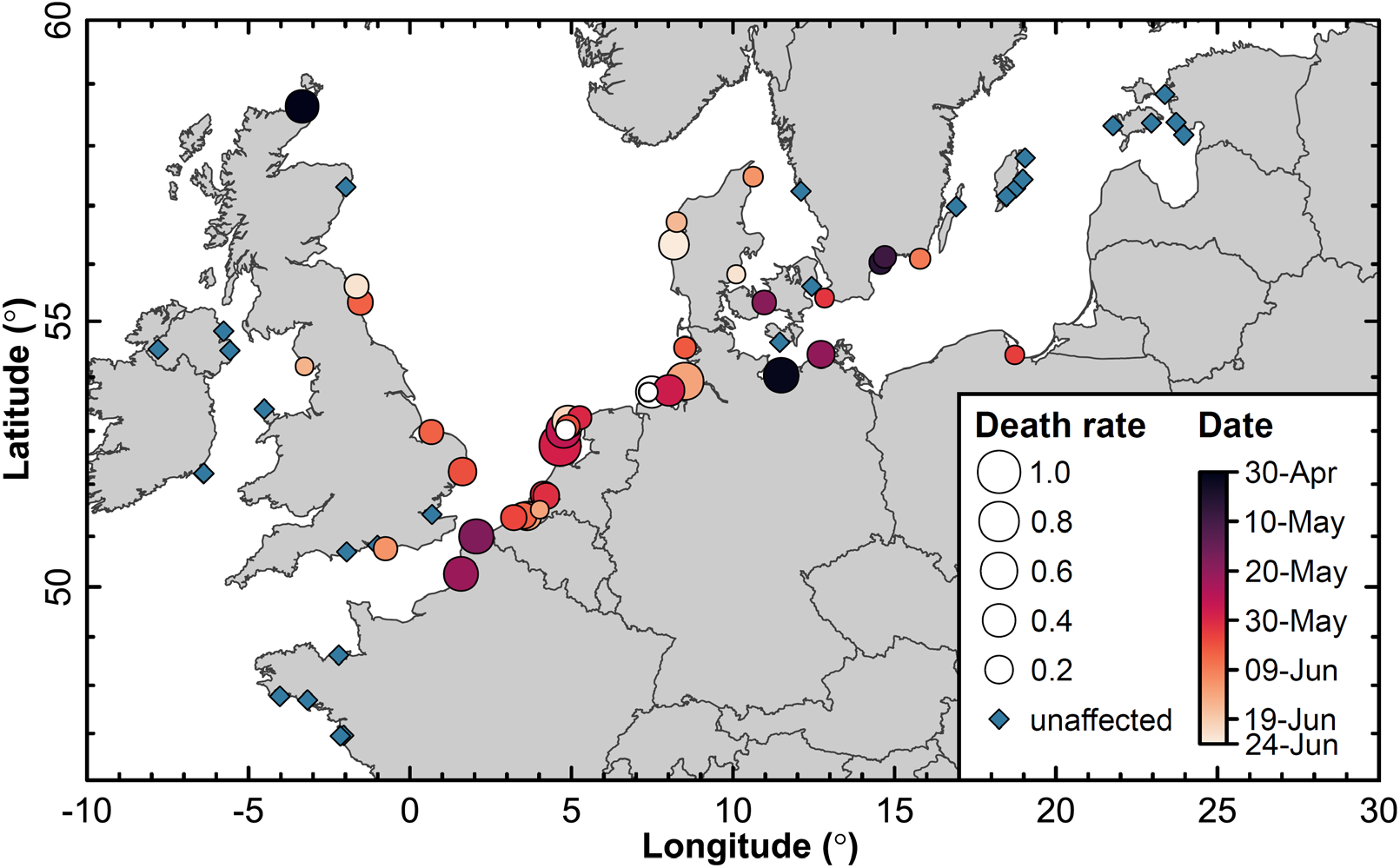
Temporal spread of HPAI in Sandwich tern colonies across northwestern Europe. Dot size reflects the severity of the outbreak (i.e., the percentage of breeding birds that were found dead) and colors represent the date of the first dead adult noticed (white: no date recorded).

Both variants circulating among Sandwich terns had previously been isolated from geese and gulls (*21, 22*). In Scotland and Germany, dead geese were found on site around two weeks prior to the outbreak in Sandwich terns. The geese in Germany tested positive (those in Scotland were not tested) and this specific variant had been previously isolated from geese (*22*). In northern France, infected herring gulls (*Larus argentatus*) were found just before the first cases of Sandwich terns. As salinity levels of the water bodies surrounding all three colonies are low, contaminated environment is a likely route of introduction of HPAI into the Sandwich tern population (*44-46*).

In May 2022, the spread of HPAI remained limited to five colonies in the vicinity of the initial two outbreaks in Germany and France. Concordantly, genetic data suggests that the eastern German variant circulated among colonies in the southern Baltic Sea for a while and only later infected colonies in northern Denmark (*22*). This was also confirmed by the temporal data, where Danish colonies were affected only early June (**Fig. 2**). From late May onwards, the spread of the virus accelerated, such that individuals in almost all Dutch, Belgian and German colonies were infected within two weeks. Phylogenetic analysis of the variants suggest that this expansion emanated from the French and Dutch outbreaks (*22*). On its march along the southern shores of the North Sea, the virus extirpated entire colonies within a week. The wave of infection subsequently reached the eastern coasts of the UK — and only receded from the northwestern European population in July (**Figs. 2 and 3**). No mortality was noted in Mediterranean breeding colonies.

**Fig. 3.**
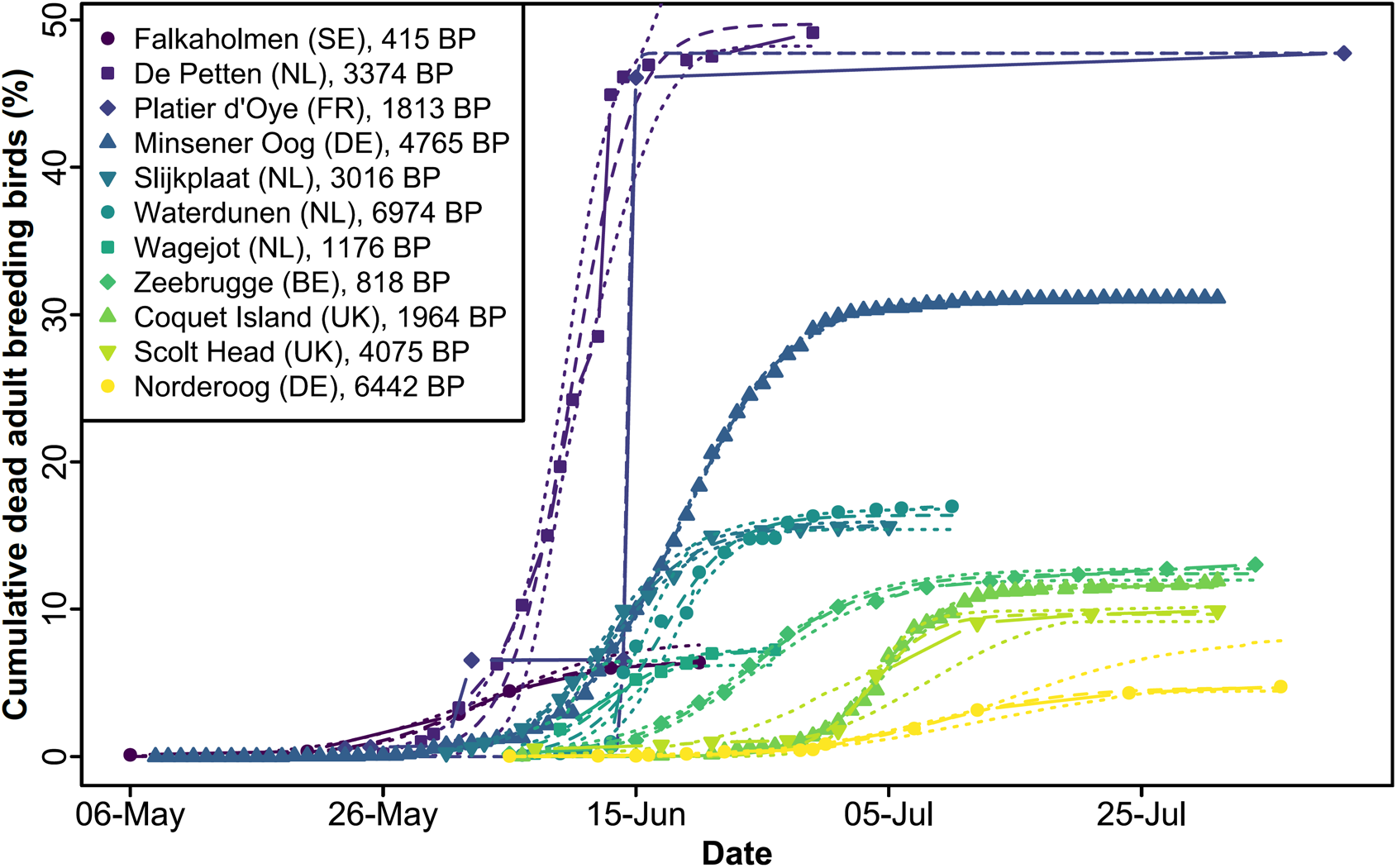
Cumulative fatality curves for eleven Sandwich tern colonies. Depicted are the raw data and parameter estimates with 95% confidence bands derived from a logistic growth model. To aid comparability between colonies of different sizes, the number of adults found dead per colony was standardized to the total number of breeding birds in a colony.

The severity of the outbreak (measured as the percentage of adult birds that were found dead within a colony) decreased with the seasonal progress (*β* = 1.46 [95% CI = 0.41–2.50], *P* = 0.006; **Figs. 2 and 4**). While it may be difficult to establish the onset of the infection retrospectively, we obtained the same result across colonies for which we had sufficient longitudinal data to estimate the inflection points and asymptotes of the fatality curves (which we used as our independent and dependent variables, respectively, *N* = 11 colonies, *P* = 0.027; **Fig. 3**). Among the many variables changing across the breeding season, increasing UV radiation and temperatures towards the summer may have limited the environmental persistence of the virus and thereby have reduced the possibility of infections through contaminated environment (*44-46*). Furthermore, the transmission rate might have been limited due to decreasing densities of birds in the colonies, because once the chicks grow older, parents lure them away from the nesting sites (*47*) into potentially less contaminated areas, which actually has been interpreted as a mechanism to avoid infectious diseases (*29*).

**Fig. 4.**
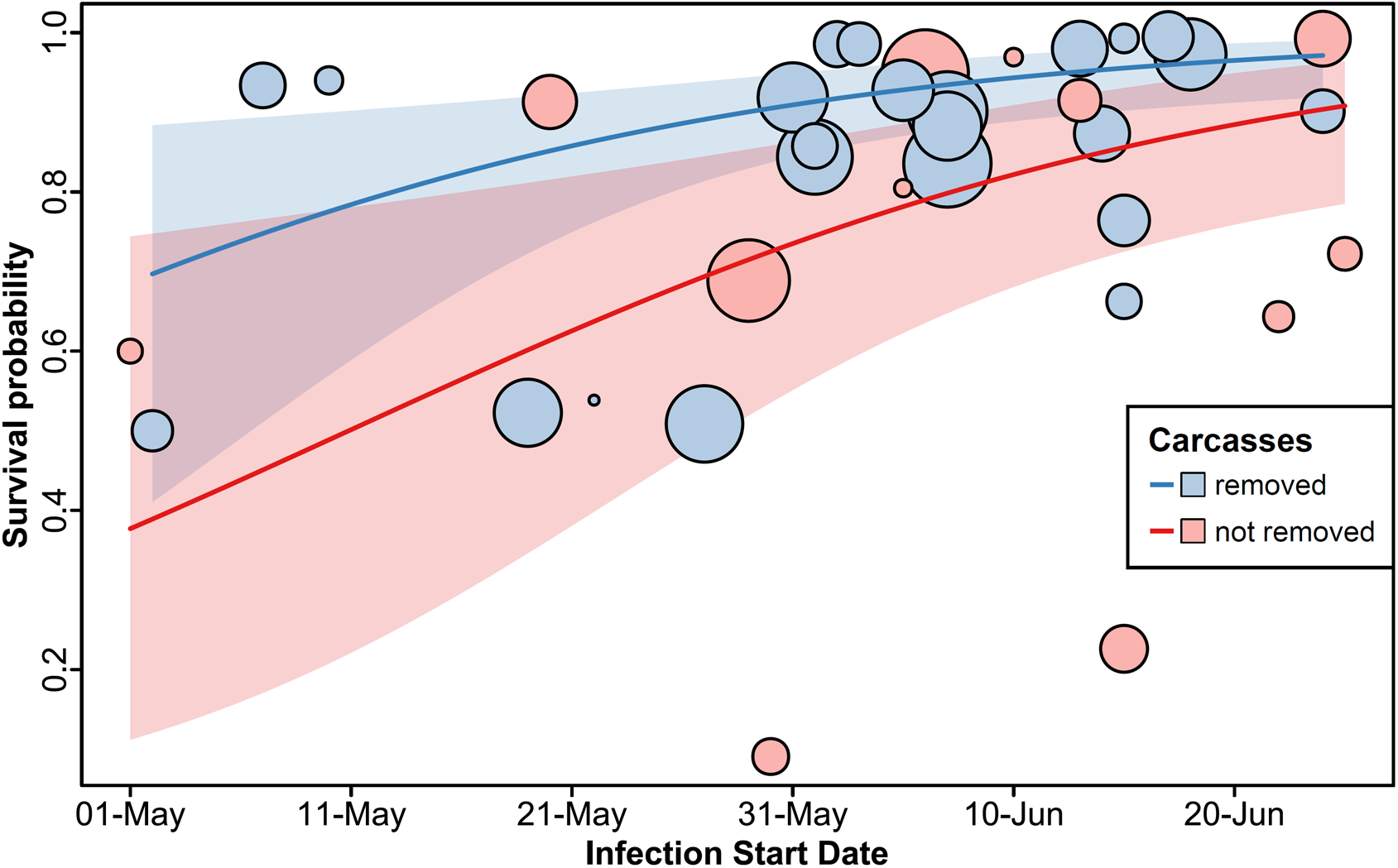
Effects of carcass removal and onset of the infection on the survival probability within an affected colony. Depicted are the raw data (colored dots) with dot size reflecting the number of breeding pairs in a colony. Estimated effects of carcass removal (no removal in red, removal in blue) and infection starting date with their 95% confidence intervals are derived from a generalized binomial linear mixed-effects model that controls for spatial autocorrelation between colonies and overdispersion.

In June, a late breeding colony was established in Belgium, which consisted of breeders from HPAI impacted and thus abandoned colonies and of younger 3–4 year-old prospecting individuals. From this colony, 35 clinically healthy adults were caught with mist nets in July. Tracheal and cloacal swab samples were collected for quantitative PCR test to detect active influenza A virus infections and thereby detect the presence of currently infected individuals that were actively shedding the virus. Blood samples were collected for serology studies to confirm pre-exposure to HPAI. Nine out of the 35 individuals tested seropositive for Asian H5 antibodies (prevalence = 25.7% [95% CI = 3.1–44.0%]). These results suggest that these birds had gained immunity from an earlier infection. The increasing presence of seropositive individuals in the population may have contributed to the decreasing mortality observed during the progress of the summer. The qPCR test targeting a highly conserved region of the viral genome, diagnostic for influenza A viruses, confirmed the absence of active influenza A virus infections in the examined individuals, meaning the tested individuals were not shedding virus anymore at the time of sampling.

All age classes (birds aged 1–27 years) were present among *N* = 403 ringed Sandwich terns found dead in Germany in 2022, forming the expected age distribution pyramid for this species (i.e. the youngest birds are largely missing because they stay at the wintering grounds in Africa; see *48*).

In 18 infected colonies monitored during the breeding season across northwestern Europe, more than 95% of all chicks died (many still in the nest). This included colonies where only a smaller fraction of breeders was found dead and adults continued to be present in the colonies. This observation suggests that HPAI easily propagated to chicks, rapidly and most likely airborne from chick to chick, and that chicks had no immunity against the virus.

In the acute situation in the 2022 breeding season, governments and local authorities applied different outbreak management approaches. In some areas carcasses were removed from Sandwich tern colonies every day or every other day, whereas at other sites, colonies were not entered and dead birds were not removed in order to reduce stress and contact between adults, chicks and humans. We used this “natural experiment” to assess carcass removal as a possible containment strategy. While controlling for the start date of the infection (see above) and spatial autocorrelation between colonies (cf. **Fig. 1**), removing carcasses reduced the severity of the outbreak in affected colonies (*β* = 1.28 [95% CI = 0.29–2.27], *P* = 0.011; **Fig. 4**). The exact transmission route of the virus from carcasses to living birds is not entirely clear, yet the viral load and infectivity remain high for prolonged times in carcasses (*49*). In 2022, Sandwich terns have been observed interacting with dead conspecifics actively, for example by pecking or copulating with them (cf. *50*), and carcass fluids may infect soil or water bodies nearby (*44, 51*).

Changes to the breeding habitat may serve as a preventive and mitigation strategy (*52*). We evaluated whether the permeability of the soil, vegetation density or the distance to fresh or brackish water had an effect on the severity of the outbreak in affected colonies. None of these explanatory variables had a significant effect (all *P* > 0.1; **Fig. S2**).

## Conclusions

The impact of the 2022 HPAI H5N1 outbreak across an entire flyway population of a colonially nesting seabird, the Sandwich tern, is unprecedented. We tally the total fatality in adult Sandwich terns at 17– 74% of the northwestern European breeding population. At least three initial entries of the H5N1 virus into the northwestern European Sandwich tern population were apparent in May 2022. Subsequently, the infection spread from these entry points and expanded through intraspecific contact within and between breeding sites. The mortality peaked in June and decreased as the breeding season advanced, possibly due to increased UV radiation and temperature in the summer (decreasing the presence of infectious viral load in the environment), decreasing density of individuals in the colonies (preventing transmission of the virus between individuals) and the increasing seroconversion of the individuals in the population. Removing carcasses from colonies was an effective containment strategy that lowered the total mortality rate by on average 15%. Given a population growth rate of 1.7% per year (*39*), it will take the northwestern European population decades to recover from the HPAI outbreak of 2022, that is, if no further outbreaks occur.

## Material and Methods

### Relevant population

European Sandwich terns are separated into three flyway populations: the Northern and Western (also referred to as “northwestern”) European population that predominantly winters along the coasts of western and southern Africa, the Mediterranean and Black Sea population which is partially synhiemic with the northwestern European population but to a large extent also stays in the Black and Mediterranean Sea during winter, and the totally allohiemic Caspian Sea population (*42, 43, 53, 54*). According to Birdlife International (*25*), the Northern and Western European population amounts to 160,000–186,000 individuals (1% = 1,700 individuals) which is 45,600–55,300 breeding pairs, the Mediterranean and Black Sea population to 62,000–221,000 individuals (1% = 1,100 individuals) and the Caspian Sea population to 110,000 individuals (1% = 1,100 individuals). We gathered data on all colonies from the Northern and Western European population, and from the closest (that is westernmost) colonies in the Mediterranean Sea, which are sometimes included in the northwestern European population. Throughout the manuscript, we refer to the Northern and Western European population excluding the Mediterranean colonies unless otherwise stated.

### Breeding data

From the 2022 breeding season, we collated data on all northwestern European Sandwich tern breeding colonies (*N* = 67 colonies with 63,116 breeding pairs, **Tab. S1**) by contacting local administrators, site managers, NGOs, scientists and stakeholders, who together form the European Sandwich Tern Research Group (see author list and acknowledgements for names and affiliations). Specifically, we requested data on the location of colonies, the number of breeding pairs and estimates of breeding success. We furthermore received data from the Mediterranean colonies in Spain (*N* = 3 colonies, 2,468 BP), southern France (*N* = 4 colonies, 3,652 BP) and Italy (*N* = 4 colonies, 2,214 BP). Some colonies in Belgium, the Netherlands, Germany, Denmark, Sweden and Ireland with in total *N* = 3,484 breeding pairs formed only late in the season, possibly by 3–4 year old birds arriving late on the breeding grounds and by birds that lost their first clutch elsewhere. Thus, we include these birds in the 63,116 BP.

### HPAI data

For each colony, we ascertained whether it was affected by HPAI. A colony was classified as being affected by HPAI if adult or juvenile Sandwich terns were dying in unusual numbers or were observed with the typical clinical signs of a HPAI infection, which include debilitation, lethargy, disorientation, loss of flight, opisthotonos, torticollis and other abnormal behaviors (*21*) that lead to death within hours after the onset of clinical signs (*N* = 39 colonies affected out of 65 colonies with HPAI status data; **Tab. S1**). We further recorded whether dead birds were tested positive for HPAI and, if so, for which virus subtype. Where a colony was classified as being affected by HPAI, we recorded the date of first fatality among adults (defined as the onset of the infection in a colony; available for *N* = 37 out of 39 affected colonies), and the total number of dead adult and juvenile Sandwich terns found. We calculated the percentage of dead adults found relative to the total number of breeding birds and used this as a measure of the severity of the HPAI outbreak. For a subset of colonies, we also collected longitudinal data to reconstruct fatality curves, which represent the cumulative number of adult Sandwich terns found dead and dying. In some countries, dead adult Sandwich terns were also counted along the shorelines (BE, NL, western DE).

There were no indications that the Mediterranean colonies were infected with HPAI and because they are often classified as belonging to a different flyway population (*53-55*), we removed those colonies from all further analyses.

### Containment and mitigation strategy data

For affected colonies, we noted whether and which containment and mitigation measures were established. The only strategy that was followed in a subset of colonies (*N* = 27 out of 37 affected colonies with known infection starting date, **Tab. S1**) was carcass removal. However, the carcass removal scheme differed between colonies. In some places, dead adults were removed every day or every other day, whereas in other colonies a week or more lay between successive carcass removal efforts. Moreover, in some colonies, not only dead adults, but also dead chicks were removed. For the final analyses, we dichotomized carcass removal, lumping all carcass removal schemes into one category. While this may seem overly simplistic, it is the most objective way of assessing the impact of carcass removal on survival probability in a colony in general. The observed effect of carcass removal, however, is likely underestimated for the following reasons: (1) It is possible that in colonies where dead adults were not collected, the total number of birds that died is underestimated, because by the end of the breeding season, when those colonies were visited and cleared from carcasses, the carcasses could have already decomposed, lost in the overgrowing vegetation or removed by predators. (2) It may be that carcass removal was only started once a colony was heavily infected, in which case we would still classify it in the carcass removal treatment group, while the positive effect of carcass removal is limited due to the late start. (3) In colonies where only dead adults were removed, dead chicks littering the colony could have been another source of infection.

### Colony-specific environmental data

Specific environmental parameters may be associated with the severity of a HPAI outbreak. We collected data on the predominant soil type in the colony (six categories), the predominant vegetation density at the beginning of June (six categories) and the distance of the colony to standing brackish or fresh water that is regularly visited by ducks, geese, gulls and Sandwich terns for washing or preening. For soil type, we established the categories (1) gravel, (2) plant material washed ashore, (3) coarse sand or very sandy soil, (4) saltmarsh or clay soil, (5) rock or concrete and (6) other soil types. These categories are ordinal with regard to water permeability. For vegetation density, we used the categories (1) bare ground, (2) plant material washed ashore, (3) patchy or sparse vegetation, (4) short vegetation, (5) in and around tussocks of long grass (e.g., lyme grass [*Leymus* sp.] or marram grass [*Ammophila* sp.]) and (6) other vegetation. These categories are ordinal with regard to vegetation density. To help people judge vegetation density and to increase interobserver agreement, we handed out drone photos of typical vegetation density categories (**Fig. S3, Tab. S2**).

### Data analyses

All statistical analyses were performed in R (v4.2.0; *56*). Spatially explicit models were fitted using the sdmTMB package (v0.1.0; *57*), which allows for the control of spatial covariation between colonies by fitting a spatial random field. To construct the random field, we used geographic coordinates in the equidistant Mercator projection and the R package INLA (v22.12.16; *58*). As Sandwich terns rarely move inland, we included the European coastline (PBSmapping R package v2.73.2; *59*) with a buffer of 10 km as a barrier in our spatial random field. Model fit was assessed via visual inspection of the distribution of residuals and using the DHARMa package (v0.4.6; *60*). We performed sensitivity analyses using different meshes with varying triangle edge length and extension distance.

We calculated the distance of each colony to the centroid of all colonies using the rgeos package (v0.6-1; *61*) and the geosphere package (v1.5-18; *62*).

We assessed whether the probability of a colony getting infected with HPAI was dependent on the size of that colony (log-transformed number of breeding pairs), the distance to the centroid of all colonies (*N* = 65 colonies with HPAI status information) and the distance to the first three outbreaks, which were most likely independent virus entry events (colonies at St John’s Pool in Scotland, at Langenwerder in Germany and at Platier d’Oye in France). To do this, we fitted a generalized linear mixed-effects model (GLMM) with a binomial error structure, taking infection status of a colony (0 = not infected, 1 = infected with HPAI, that means without considering colony size) as the dependent variable, and colony size, distance to the center and distance to first infections as three covariates. The distance to first infections showed a trend in the expected direction (being closer increased the probability of a colony getting infected; *β* = 2.41 [95% CI = -0.80–5.62], *P* = 0.14).

Next, we tested whether the severity of a HPAI outbreak was explained by the onset of the infection in a colony and by the carcass removal containment strategy. For that, we restricted the dataset to those colonies with a HPAI infection (*N* = 37 colonies with known infection starting date and carcass removal information). We fitted a spatially explicit binomial GLMM, using the number of birds not found dead divided by twice the number of breeding pairs as our dependent variable (which we took as the survival probability). In order to incorporate differences in colony size, we weighted the dependent variable by the number of breeding birds (= 2 × BP). We fitted the date of the first case detected as a covariate and carcass removal as a factor with two levels (yes or no) as our predictors. We transformed both predictors using the rescale function from the arm package (v1.13-1; *63*), which sets the mean to 0 and divides by two standard deviations to ensure comparability (cf. *64*). We furthermore fitted an observation-level random effect to account for overdispersion.

Lastly, we assessed whether the severity of a HPAI outbreak was associated with colony-specific environmental variables. To do this, we again restricted the data set to those 37 colonies in which HPAI infection was confirmed and the starting date of the outbreak was documented. We fitted a spatially explicit binomial GLMM with survival probability as our dependent variable, weighted by the number of breeding birds. Because the onset of the infection and carcass removal were significantly associated with survival probability, we fitted these two predictors and added either soil type (five levels) or vegetation density (four levels) as further explanatory variables. From the initially anticipated six categories for each of these predictors (see above) some were not observed. In separate models, we fitted soil type and vegetation density as covariates (using one degree of freedom each). Including an observation-level random effect prevented the model from converging, so this was dropped from these models.

Because the onset of an infection may be mis specified when a colony is not visited on the day of the first HPAI fatality case, we collected longitudinal data in 11 colonies to fit logistic curves to the cumulative number of dead adults collected in a colony over time. For each colony, we estimated the asymptotes and the inflection points of the curves using nonlinear least-squares regression. We used these two variables in a subsequent linear model, to assess whether the asymptotes (which represent the severity of the outbreaks) depended on the inflection points (i.e., when half the birds had died).

In order to predict the number of adult Sandwich terns dying outside the colonies, we fitted a major axis regression using the R package smatr (v3.4-8; *65*). For a subset of countries, we obtained data on the number of Sandwich terns found dead outside the colonies. We used these data as our dependent variable, and the number of birds found dead inside the colonies as our sole predictor. Subsequently, for all countries, we predicted the number of birds found dead outside the colonies from the number of dead birds inside the colonies.

### Sero-surveillance and monitoring of HPAI infections in a late-established sandwich tern colony

This surveillance was part of the wildlife HPAI disease surveillance of the Flemish government in Belgium. A late breeding colony was established near Zeebrugge (BE) in June, which consisted of breeders from HPAI impacted and thus abandoned colonies and of younger 3–4 year-old prospecting individuals. On July 18^th^, when birds were still raising chicks at this colony, 35 adults were caught there with mist nets at night on the adjacent beach. Consequently, our population sample might have included some birds that were prospecting, dispersing or used Zeebrugge as a stopover during migration. This was confirmed by sightings of color rings and head molt patterns (*66*). Approximately 300 μl of blood was taken from the brachial vein by a trained veterinarian and subsequently the serum was separated from clotted blood by centrifugation (1227 g for 5 min). Sera were screened by the AI National Reference Laboratory of Belgium for influenza A nucleoprotein specific antibodies using the ID Screen^®^ NP competition enzyme-linked immunosorbent assay (ELISA; IDVet) according to the manufacturer’s guidelines. The NP-ELISA test is considered positive with competition <45%. NP-ELISA positive samples were subjected to hemagglutination inhibition (HAI) testing with different sets of H5-antigens, which allows the differentiation of Eurasian H5-from an Asian GsGd-clade H5-immune response. The HAI test conformed to (*67*). Briefly, reference antigens were diluted at four HA-units and tested with a 0.5 serial dilution of sera under investigation. HAI-values > 3 × log^2^(1/8) were considered as positives. Sensitivity and specificity of NP-ELSIA have not been ascertained in wild birds and vary among chickens and ducks (chickens: 95% CI sensitivity of 0.72–1.00 and 95% CI specificity of 0.91–1.00, ducks: 95% CI sensitivity of 0.72–1.00 and 95% CI specificity of 0.72–1.00). We here use the widest CI ranges to obtain confidence intervals (see below).

For the same 35 birds, the presence of avian influenza virus shed via the oral/cloacal route was assessed by the AI National Reference Laboratory of Belgium. Oropharyngeal and cloacal swabs were collected from each bird and were immediately placed in viral transport medium and analysed in pools of a maximum of five individuals. Briefly, viral RNA was extracted from 200 µl swab fluid with the semi-automated Indimag Pathogen Kit^®^ on the IndiMag 48s (Indical). Viral RNA detection was performed by TaqMan^®^ real-time (RT)-PCR using a universal oligoset, which detects the highly conserved M-gene, diagnostic for influenza A viruses (*67, 68*). The AgPath-ID™ One-Step RT-PCR kit (Thermo Fisher Scientific) was used for amplification of the viral RNA according to the manufacturer’s instructions on the LightCycler^®^ 480 RT-PCR system (Roche). Sample quality and extraction efficacy were evaluated by the detection of the beta-actine endogenous control (cf. *69*).

We analyzed the sero-surveillance data using sensitivity and specificity values asserted in ducks, and estimated the prevalence of seropositive individuals with 95% confidence intervals using the bootComb R package (v1.1.2; *70*).

### Surveys at major roosts

Sandwich terns show post-breeding dispersal before migrating to their wintering grounds along the western and southern African coasts (*42*). Ring recovery data indicate that birds breeding along the shores of the southern North Sea (but partly also around the Baltic Sea and rarely in Great Britain) gather in Danish North Sea waters in late summer. It has been estimated that 10,000 birds roost on the island Fanø alone every year (*71*), because of high local prey abundance. This makes Fanø a representative sample of birds breeding along the shores of the southern North Sea, which were most heavily affected by HPAI. From 2020–2022, the number of adults and first-year birds (that is juvenile birds hatched in the respective year) roosting on Fanø were recorded between July and September.

The percentage of first-year birds at major northwestern European roosts in Denmark was analyzed with binomial GLMMs in the R package lme4 (v1.1-32; *72*). The proportion of first-year birds was fitted as the dependent variable, weighted by the number of birds observed. Scaled date was fitted as a quadratic and linear covariate (using the poly() R function) in interaction with year (factor with three levels: 2020, 2021, 2022). Location was included as a random intercept. Confidence intervals, marginal effects and contrasts were estimated using the ggeffects package (v1.2.1; *73*).

## Acknowledgements

We are enormously grateful to all people and organizations contributing data on the HPAI outbreak in the Sandwich tern breeding season 2022: Antoine Arnaud, Thomas Blanchon, Marco Basso, Amy Burns, Bernard Cadiou, Philippe Carruette, Olivier Enjalbert, Jitske Esselaar, Duncan Halpin, Robin Harvey, Bernd Heinze, Mikael Hellman, Yann Jacob, Luke Johns, Rebecca Jones, Rémi Jullian, Alain Le Dreff, Meelis Leivits, Adeline Leray, Mhairi Maclauchlan, Murray Ochard, Parc Natural del Delta de l’Ebre (Departament de Acció Climàtica, Alimentació y Acció Rural, Generalitat de Catalunya), Patrice Peron, Jean-Roger Perrot, Kathleen Perrot, Laurie Pescayre, Dominique Robart, Carolin Rothfuß, Vincent Rotureau, Philippe Sauvaget, Daryl Short, Alexandre Sibille, Hugh Thurgate, Roberto Tinarelli, Rémi Tiné, Jens Umland, Marc Van de Walle, Nicolas Vanermen, Hilbran Verstraete, Guillaume Villette, Izzy Williamson, Chris Wynne, Barry Yates. Jan Baert, Lowie Lams and Annemarie Waerendorff assisted for the serology sampling and Sciensano (Federal Research Institute for Public and Animal Health in Belgium) kindly conducted serological analyses. Sean Anderson provided invaluable support in sdmTMB. We further thank the Common Wadden Sea Secretariat for organizing workshops on avian influenza in colonial seabirds. This work was funded by the Common Wadden Sea Secretariat to UK; the Ministry of Agriculture, Nature and Food Quality of the Netherlands to ML; OFB, DREAL Occitanie, DREAL PACA to OS; EU LIFE (LIFE on the edge: LIFE19NAT/UK/964) to WS; and the Ministry of Infrastructure and Water Management of the Netherlands to RvB.

## Ethics statement

Blood samples were taken under the wildlife HPAI disease surveillance of the Flemish government in Belgium.

## Data availability

Data and analyses scripts are available in the Supplement and through the Open Science Framework (doi: https://osf.io/y89k6/?view_only=81b6fa5ca42a4a849bb4e0d896a25bff).

## Author contributions

IA, MB, TB, AB, SB, AC, WC, JD, EE, RF, BH, MH, VH, CH, RtV, EK, UK, MK, SK, KL, RL, NL, ML, SL, LL, RM, HM, WM, PM, SN, PO, FP, KTP, CR, FSc, FSe, OS, LS, JSm, WS, JSt, ES, VS, RGV, RvB, JV, EW, MW collected HPAI data in the field. MV performed serological tests. KF collected data on Danish roosts. UK, WC and TB designed the study. UK summarized and analyzed the data with input from WC. UK and WC wrote the first draft of the manuscript and prepared the final manuscript with input from all authors. All authors approved the final manuscript.

