## Supplement for "Highly pathogenic avian influenza causes mass mortality in Sandwich tern (*Thalasseus sandvicensis*) breeding colonies across northwestern Europe"

[13] 24241 Blumenthal, Germany

[14] University of Hamburg, Institute of Cell and Systems Biology of Animals, Animal Ecology and Conservation, 20146 Hamburg, Germany

[15] Landesamt für Umwelt, Naturschutz und Geologie MV, 18273 Güstrow, Germany

[16] Staatsbosbeheer Zuid-Hollandse Delta, Numansdorp, the Netherlands

[17] Falsterbo Bird Observatory, Sweden

[18] Wageningen Marine Research, 1781AG Den Helder, the Netherlands

[19] Kalmar Maritime Academy, Linnaeus University, 391 82 Kalmar, Sweden

[20] Torhamn Bird Observatory, Sweden

[21] Natural England, UK

[22] Communauté de Communes de l'île de Noirmoutier, Noirmoutier-en-l'île, France

[23] Terrestrial Ecology Unit, Department of Biology, Ghent University, Ghent, Belgium

[24] Department of Vertebrate Ecology & Zoology, University of Gdańsk, Gdańsk, Poland

[25] Coast Zone Conservation, UK

[26] Birdwatch Ireland, Kilcoole, Ireland

[27] 31196 Heberg, Sweden

[28] Lower Saxon Wadden Sea National Park Authority, 26382 Wilhelmshaven, Germany

[29] DK-2650 Hvidovre, Denmark

[30] School of Natural and Environmental Sciences, Newcastle University, Newcastle upon Tyne, UK

[31] SELC Società Cooperativa, Marghera Venezia, Italy

[32] Stichting Het Zeeuwse Landschap, Wilhelminadorp, the Netherlands

[33] Conservatoire d'espaces naturels d'Occitanie, 26 allées de Mycènes, 34000 Montpellier, France

[34] Area per l'Avifauna Migratrice (BIO-AVM) - Istituto Superiore per la Protezione e la Ricerca Ambientale (ISPRA), Ozzano dell'Emilia BO, Italy

[35] St. John's, Brough, Thurso, Caithness, KW14 8YD, Scotland, UK

[36] Research Institute for Nature and Forest, 1000 Brussels, Belgium

[37] 30125 Venice, Italy

[38] 7345 CC Wenum-Wiesel, the Netherlands

[39] Agency for Nature and Forests, 1000 Brussels, Belgium

[40] Comers Wood Croft, Aberdeenshire, UK AB51 7QB, UK

### INDEX


| | |
|---|---|
| Supplementary Text | 4 |
| Supplementary Figures | 6 |
| Supplementary Tables | 9 |
| References | 13 |

#### SUPPLEMENTARY TEXT

*Recommendations to breeding site managers*

There are no national or international structures in place that record the total numbers of wild birds dying due to an infection with avian influenza, as this disease is still treated mainly as an economic, agricultural and human-health related problem, rather than a severe threat to protected wildlife (*1*). Thus, an active monitoring and surveillance strategy of the virus and associated mortality in wild birds should be implemented, allowing for a rapid intervention once HPAI is introduced into a Sandwich tern (or other seabird species) breeding colony.

Removing carcasses seems to be an effective containment strategy that should at least be implemented in those colonies that are looked after intensively. In order to be most effective, removal should start as early as possible to keep disturbances at a minimum and to reduce the risk of infection from carcasses. Removal schemes should be fully documented (including records of the number and age of the birds found dead and metal- and color-ring codes). The advantages and disadvantages of carcass removal are reviewed in (*2, 3*). At certain phases of the breeding cycle, entering and disturbing a colony for prolonged periods of times can lead to increased aggression between birds, reduced breeding success and it can cause abandonment of entire colonies. Thus, potentially infected birds are forced to disperse and may rebreed elsewhere (*4*), which could spread HPAI further to breeding sites that had not been affected before. As a consequence, breeding colonies must be intensively monitored to detect any early signs of HPAI and access must be granted to those people and organizations looking after the breeding sites, even if HPAI is to return. It is of vital importance for people working in the colonies to wear adequate personal protective equipment (PPE) to minimize the risk of viral spillover to humans. Only people vaccinated against human flu should work in bird flu infected colonies, to reduce the risk of a recombination event between the mammalian and avian influenza virus. Furthermore, the human influenza vaccination also appears to promote restricted protection against avian influenza (*5*). A detailed risk assessment is currently conducted by an expert group under the direction of the Common Wadden Sea Secretariat and the Friedrich-Loeffler-Institute. Furthermore, an expert group under the direction of the Common Wadden Sea Secretariat has developed guidelines for mitigation and data collection for avian influenza in bird colonies in the Wadden Sea (*3*).

*Recommendations for further studies*

For a better understanding of why Sandwich terns were particularly susceptible to HPAI, more research on the sources and modes of infection, incubation times, effective containment and immunity is urgently needed. This includes (i) population ecological studies on the exchange of Sandwich terns between breeding sites (cf. *4, 6*) and (ii) detailed survival analyses and demographic modelling using ring recovery data (cf. *7*). Thus, the research on the natural history of this species should be extended, which means breeding colonies should remain accessible to scientists. (iii) The extent of survival and immunity should be assessed in larger samples and more detail in live birds captured at the breeding grounds. (iv) Containment strategies (for example carcass removal) should be documented well (for example including experimental designs) to enable a better evaluation of the effects of these measures.

#### Supplementary Figures

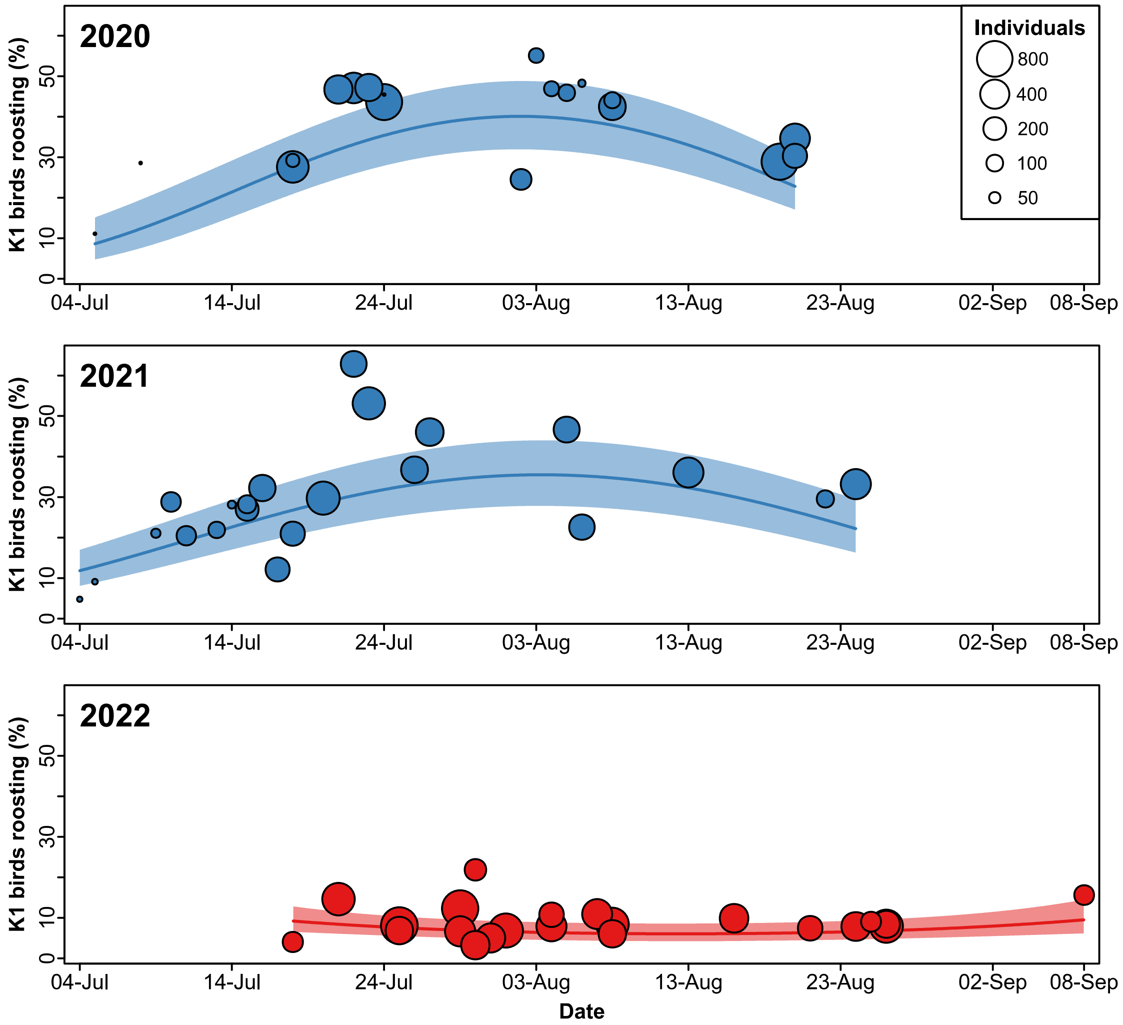


**Fig. S1** | Percentage of first-year birds observed at the major roost sites on the island Fanø in the Danish part of the Wadden Sea in the years 2020, 2021 and 2022. There was a significant interaction between year and date fitted as a quadratic effect. Displayed are the regression lines with 95% confidence intervals (CIs). The estimated marginal percentage of first-year birds at the end of July (30th/31st, which is the mean date) was 39.7 (95% CI = 31.6–48.3) in 2020, 34.9 (95% CI = 27.4–43.3) in 2021 and 6.8 (95% CI = 4.9–9.3) in 2022.

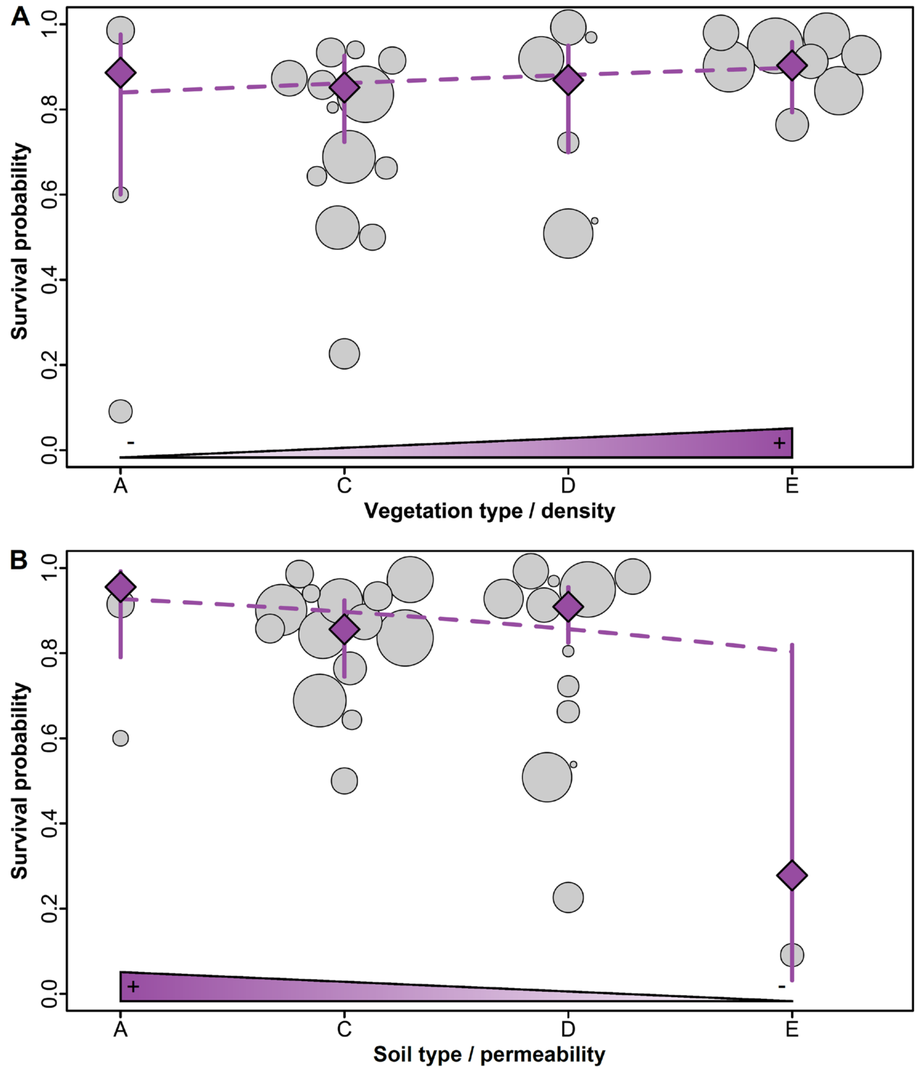


**Fig. S2 |** The effect of environmental factors on survival probability across colonies. (A) Vegetation type / density and (B) soil type / permeability. Vegetation and soil categories A–E are ordinal in respect to density and permeability. For vegetation density, category A represents bare ground and category E represents in and around tussocks of long grass (cf. Fig. S2). For soil type, category A represents gravel and category E rock or concrete. We fitted vegetation and soil type either as a factor with four levels (depicted are parameter estimates with 95% confidence intervals in purple) or as a covariate using one degree of freedom (purple broken line, $P$ = 0.46 for vegetation and $P$ = 0.17 for soil) in a spatially explicit mixed effects model.

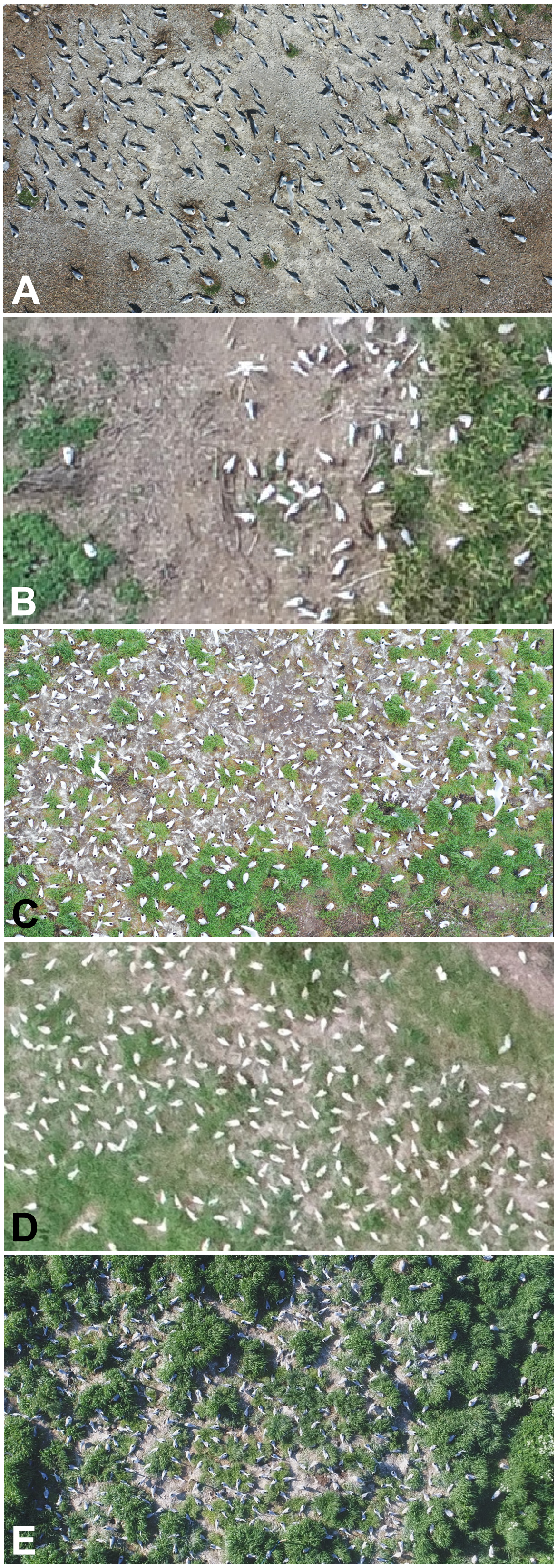


**Fig. S3 |** Five categories of vegetation type / density. (A) bare ground, (B) plant material washed ashore, (C) patchy or sparse vegetation, (D) short vegetation, (E) in and around tussocks of long grass (e.g., lyme grass [*Leymus* sp.] or marram grass [*Ammophila* sp.]). In 2022, plant material washed ashore was not reported from any colony.

#### Supplementary Tables

**Tab. S1 |** Breeding and HPAI data collated on all northwestern European and the westernmost Mediterranean Sandwich tern breeding colonies.

| Country | Colony | Latitude | Longitude | *N* breeding pairs | Affected | Late colony | Infection start date | *N* dead adults | Dead adults removed |
|---|---|---|---|---|---|---|---|---|---|
| ES | Punta de la Banya 1 | 40.5823 | 0.6990 | 585 | FALSE | FALSE | | 0 | |
| ES | Punta de la Banya 2 | 40.5844 | 0.7035 | 1609 | FALSE | FALSE | | 0 | |
| ES | Punta del Fangar | 40.7889 | 0.7640 | 274 | FALSE | FALSE | | 0 | |
| IT | Lagoon of Venice – South | 45.2530 | 12.2344 | 1469 | FALSE | FALSE | | 0 | |
| IT | Lagoon of Venice – North | 45.4410 | 12.3565 | 150 | FALSE | FALSE | | 0 | |
| IT | Pialassa dei Piomboni, Po Delta | 44.4594 | 12.2778 | 560 | FALSE | FALSE | | 0 | |
| IT | Sacca di Scardovari, Po Delta | 44.8603 | 12.4178 | 35 | FALSE | FALSE | | 0 | |
| FR | A l'est de l'Herault | 43.6117 | 4.0969 | 244 | FALSE | FALSE | | 0 | |
| FR | Camargue | 43.4699 | 4.2809 | 465 | FALSE | FALSE | | 0 | |
| FR | Île aux Moutons | 47.7755 | -4.0292 | 453 | FALSE | FALSE | | 0 | |
| FR | Ile de la Colombière | 48.6235 | -2.2061 | 291 | FALSE | FALSE | | 0 | |
| FR | Iniz er mour | 47.6983 | -3.1734 | 502 | FALSE | FALSE | | 0 | |
| FR | Lagune de Bouin | 46.9562 | -2.0509 | 130 | FALSE | FALSE | | 0 | |
| FR | Parc du Marquenterre Bay of Somme | 50.2575 | 1.5787 | 35–44 | TRUE | FALSE | 21-May | 36 | TRUE |
| FR | Platier d'Oye National Nature Reserve | 50.9940 | 2.0535 | 1813 | TRUE | FALSE | 18-May | 1731 | TRUE |
| FR | Polder de Sébastopol, Noirmoutier | 46.9431 | -2.1547 | 4400 | FALSE | FALSE | | 0 | |
| FR | Salin de Berre | 43.4803 | 5.1341 | 2 | FALSE | FALSE | | 0 | |
| FR | Sète | 43.3799 | 3.6179 | 2941 | FALSE | FALSE | | 0 | |
| BE | Sternenschiereiland, Zeebrugge/Heist | 51.3599 | 3.2136 | 13 | TRUE | FALSE | 03-Jun | 11 | TRUE |
| BE | Sternenschiereiland, Zeebrugge/Heist | 51.3599 | 3.2136 | 805 | TRUE | TRUE | 13-Jun | 204 | TRUE |
| NL | Bliek | 51.7765 | 4.2171 | 404 | TRUE | FALSE | 31-May | 115 | TRUE |
| NL | De Petten | 53.0117 | 4.7554 | 3374 | TRUE | FALSE | 26-May | 3316 | TRUE |
| NL | De Putten | 52.7371 | 4.6470 | 220 | TRUE | FALSE | 29-May | 400 | FALSE |
| NL | De Steenplaat | 53.1826 | 4.8940 | 150 | TRUE | TRUE | 21-Jun | 107 | FALSE |
| NL | Griend | 53.2519 | 5.2541 | 2100 | TRUE | FALSE | 30-May | 342 | TRUE |

| | | | | | | | | | |
|---|---|---|---|---|---|---|---|---|---|
| NL | Hooge Platen | 51.3918 | 3.6202 | 600 | TRUE | FALSE | 14-Jun | 283 | TRUE |
| NL | Koude- en Kaarspolder | 51.5156 | 4.0175 | 137 | TRUE | FALSE | 14-Jun | 2 | TRUE |
| NL | Slijkplaat | 51.8009 | 4.1433 | 3016 | TRUE | FALSE | 31-May | 941 | TRUE |
| NL | Wagejot | 53.0895 | 4.8979 | 1176 | TRUE | FALSE | 04-Jun | 170 | TRUE |
| NL | Prins Hendrik Zanddijk | 53.0231 | 4.8199 | 600 | TRUE | TRUE | NA | 30 | TRUE |
| NL | Waterdunen | 51.3978 | 3.5094 | 6974 | TRUE | FALSE | 06-Jun | 2292 | TRUE |
| DE | Baltrum (FK609) | 53.7229 | 7.3740 | 208 | TRUE | FALSE | NA | 6 | FALSE |
| DE | Baltrum (FK610 late) | 53.7205 | 7.4003 | 50 | NA | TRUE | NA | NA | FALSE |
| DE | Baltrum (FK610) | 53.7231 | 7.3831 | 391 | TRUE | FALSE | NA | 8 | FALSE |
| DE | Barther Oie | 54.4087 | 12.7277 | 11 | TRUE | FALSE | 20-May | 4 | TRUE |
| DE | Langenwerder | 54.0277 | 11.4930 | 300 | TRUE | FALSE | 01-May | 300 | TRUE |
| DE | Langeoog | 53.7260 | 7.4840 | 165 | TRUE | TRUE | NA | 112 | FALSE |
| DE | Minsener Oog | 53.7567 | 8.0214 | 4765 | TRUE | FALSE | 28-May | 2967 | FALSE |
| DE | Neuwerk | 53.9181 | 8.5112 | 462 | TRUE | FALSE | 14-Jun | 715 | FALSE |
| DE | Neuwerk | 53.9305 | 8.4970 | 200 | TRUE | TRUE | 14-Jun | 135 | TRUE |
| DE | Norderoog | 54.5290 | 8.5157 | 6442 | TRUE | FALSE | 05-Jun | 650 | FALSE |
| DK | Agger Tange | 56.7233 | 8.2550 | 2400 | TRUE | FALSE | 17-Jun | 131 | TRUE |
| DK | Fjandø | 56.3382 | 8.1728 | 180 | TRUE | TRUE | 24-Jun | 100 | FALSE |
| DK | Hirsholm | 57.4854 | 10.6236 | 796 | TRUE | FALSE | 12-Jun | 32 | TRUE |
| DK | Hjarnø | 55.8252 | 10.1010 | 789 | TRUE | FALSE | 23-Jun | 12 | FALSE |
| DK | Sprogø | 55.3324 | 10.9661 | 712 | TRUE | FALSE | 19-May | 124 | FALSE |
| DK | Saksfjed Inddæmning | 54.6268 | 11.4494 | 8 | FALSE | TRUE | | 0 | |
| DK | Holmesø, Køge Bugt | 55.6142 | 12.4382 | 45 | FALSE | FALSE | | 0 | |
| PL | Gdańsk | 54.3980 | 18.7242 | 350 | TRUE | FALSE | 02-Jun | 10 | TRUE |
| SE | Eneskärskläpparna, Karlskrona | 56.0951 | 15.7933 | 30–100 | TRUE | FALSE | 09-Jun | 4 | FALSE |
| SE | Falkaholmen, Blekinge | 56.0297 | 14.5712 | 400 | TRUE | FALSE | 06-May | 53 | TRUE |
| SE | Falkaholmen, Blekinge | 56.0297 | 14.5712 | 15 | TRUE | TRUE | NA | 0 | FALSE |
| SE | Gräsholmen, Gotland | 57.2936 | 18.7522 | 61 | FALSE | FALSE | | 0 | |
| SE | Gräsholmen, Gotland | 57.2936 | 18.7522 | 15 | FALSE | TRUE | | 0 | |
| SE | Landgrens Holme, Skåne | 55.4128 | 12.8370 | 415 | TRUE | FALSE | 01-Jun | 12 | TRUE |
| SE | Landgrens Holme, Skåne | 55.4128 | 12.8370 | 41 | FALSE | TRUE | | 0 | |

| | | | | | | | | | |
|---|---|---|---|---|---|---|---|---|---|
| SE | Norrören, Blekinge | 56.1252 | 14.7015 | 125 | TRUE | FALSE | 09-May | 15 | TRUE |
| SE | Östergarnsholm, Gotland | 57.4411 | 18.9756 | 37 | FALSE | FALSE | | 0 | |
| SE | Sigdesholmen, Gotland | 57.1539 | 18.4658 | 210 | FALSE | FALSE | | 0 | |
| SE | Sigdesholmen, Gotland | 57.1539 | 18.4658 | 25 | FALSE | TRUE | | 0 | |
| SE | Skenholmen, Gotland | 57.7969 | 19.0431 | 87 | FALSE | FALSE | | 0 | |
| SE | Tärnön, Bua, Halland | 57.2424 | 12.1063 | 70 | FALSE | FALSE | | 0 | |
| SE | Villgrund, Öland | 56.9850 | 16.9120 | 87 | FALSE | FALSE | | 0 | |
| GB | Brownsea Island Poole Harbour | 50.6931 | -1.9607 | 227 | FALSE | FALSE | | 0 | |
| GB | Cemlyn, Wales | 53.4100 | -4.5130 | >2000 | FALSE | FALSE | | 0 | |
| GB | Coquet Island | 55.3350 | -1.5384 | 1964 | TRUE | FALSE | 06-Jun | 461 | TRUE |
| GB | Forvie NNR, North-east Scotland | 57.3137 | -1.9860 | 1031 | FALSE | FALSE | | 0 | |
| GB | Gravel Ridge Island ASSI, Northern Ireland | 54.5013 | -7.7994 | 102 | Unknown | FALSE | NA | More than usual | NA |
| GB | Hodbarrow, Cumbria | 54.1919 | -3.2663 | 580 | TRUE | FALSE | 16-Jun | 6 | TRUE |
| GB | Inner Farne, Farne Islands | 55.6166 | -1.6549 | 350 | TRUE | FALSE | 23-Jun | 69 | TRUE |
| GB | Langstone Harbour | 50.8233 | -1.0069 | 9 | FALSE | FALSE | | 0 | |
| GB | Larne Lough, Northern Ireland | 54.8229 | -5.7714 | 1254 | FALSE | FALSE | | 0 | |
| GB | Medway Islands | 51.4205 | 0.6805 | 200 | FALSE | FALSE | | 0 | |
| GB | Pagham Harbour | 50.7563 | -0.7594 | 335 | TRUE | FALSE | 12-Jun | 57 | FALSE |
| GB | RSPB Minsmere Nature Reserve | 52.2409 | 1.6261 | 64 | TRUE | FALSE | 04-Jun | 25 | FALSE |
| GB | Scolt Head Norfolk | 52.9855 | 0.6609 | 4075 | TRUE | FALSE | 06-Jun | 805 | TRUE |
| GB | St John's Pool, Caithness, Scotland | 58.6370 | -3.3410 | 100 | TRUE | FALSE | 30-Apr | 80 | FALSE |
| GB | Strangford Lough, Northern Ireland | 54.4773 | -5.5770 | 400 | FALSE | FALSE | | 0 | |
| IE | Inish, Lady’s Island Lake, Wexford | 52.2023 | -6.3908 | 824 | FALSE | FALSE | | 0 | |
| IE | Inish, Lady’s Island Lake, Wexford | 52.2023 | -6.3908 | 110 | FALSE | FALSE | | 0 | |
| IE | Sgarbheen, Lady’s Island Lake, Wexford | 52.2091 | -6.3854 | 726 | FALSE | FALSE | | 0 | |
| IE | Sgarbheen, Lady’s Island Lake, Wexford | 52.2091 | -6.3854 | 76 | FALSE | FALSE | | 0 | |
| EE | Anemaa, Matsalu | 58.8290 | 23.3725 | 80 | FALSE | FALSE | | 0 | |
| EE | Nootamaa, Vilsandi Rahvuspark | 58.3231 | 21.7665 | 14 | FALSE | FALSE | | 0 | |
| EE | Laid nr. 10, Kunnati laht | 58.3696 | 22.9557 | 632 | FALSE | FALSE | | 0 | |
| EE | Soondrerank, Matsi rand | 58.3791 | 23.7212 | 420 | FALSE | FALSE | | 0 | |
| EE | Umalaid, Kihnu | 58.1723 | 23.9586 | 469 | FALSE | FALSE | | 0 | |

**Tab. S2 |** Environmental data collated on HPAI positive Sandwich tern colonies.

| Country | Colony | Distance to standing water (km) | Soil type* | Vegetation density† | Remarks |
|---|---|---|---|---|---|
| FR | Parc du Marquenterre Bay of Somme | 0.06 | D | D | |
| FR | Platier d'Oye National Nature Reserve | 0 | F | C | 3F. Sand and silt (limon) |
| BE | Sternenschiereiland, Zeebrugge/Heist | 0.05 | C | C | |
| NL | Bliek | 0.05 | C | C | |
| NL | De Petten | 0.01 | D | D | |
| NL | De Putten | 0.01 | E | A | |
| NL | De Steenplaat | 2.0 | C | C | |
| NL | Griend | 0.15 | C | D | |
| NL | Hooge Platen | 0.05 | C | E | |
| NL | Koude- en Kaarspolder | | | | |
| NL | Slijkplaat | 0.05 | C | E | |
| NL | Wagejot | 0.01 | D | E | |
| NL | Prins Hendrik Zanddijk | 0.8 | C | C | |
| NL | Waterdunen | 0.03 | C | C | |
| DE | Baltrum (FK609) | None | D | E | |
| DE | Baltrum (FK610) | None | D | E | |
| DE | Barther Oie | None | F | D | Salt marsh peat |
| DE | Langenwerder | None | C | C | |
| DE | Langeoog | None | C | C | |
| DE | Minsener Oog | None | C | C | |
| DE | Neuwerk | 0.02 | D | C | |
| DE | Neuwerk | 0.02 | D | C | |
| DE | Norderoog | None | D | E | |
| DK | Agger Tange | 0.4 | C | E | |
| DK | Fjandø | None | D | D | |
| DK | Hirsholm | 1.2 | D | E | |
| DK | Hjarnø | 0 | D | D | |
| DK | Sprogø | 0.2 | D | E | |
| PL | Gdańsk | 7.0 | C | A | |
| SE | Eneskärskläpparna, Karlskrona | 8.0 | D | D | Peat and plant material |
| SE | Falkaholmen, Blekinge | 1.0 | C | C | |
| SE | Landgrens Holme, Skåne | | | | |
| SE | Norrören, Blekinge | 1.0 | C | C | |
| GB | Coquet Island | | | | |
| GB | Hodbarrow, Cumbria | | | | |
| GB | Inner Farne, Farne Islands | | | | |
| GB | Pagham Harbour | 1 | A | C | |
| GB | RSPB Minsmere Nature Reserve | 0 | D | C | |
| GB | Rye Harbour | 0.1 | A | C | |
| GB | Scolt Head Norfolk | 2 | C | E | |
| GB | St John's Pool, Caithness, Scotland | 0 | A | A | |

* (A) gravel, (B) plant material washed ashore, (C) sand or very sandy soil, (D) saltmarsh or clay soil, (E) rock or concrete and (F) other soil types.

† (A) bare ground, (B) plant material washed ashore, (C) patchy or sparse vegetation, (D) short vegetation, (E) in and around tussocks of long grass (e.g., lyme grass [*Leymus* sp.] or marram grass [*Ammophila* sp.]) and (F) other vegetation.
